# Deep Brain Stimulation of Nucleus Basalis of Meynert improves learning in rat model of dementia

**DOI:** 10.1101/2024.04.05.588271

**Authors:** Deepak Kumbhare, Megan Rajagopal, Jamie Toms, Anne Freelin, George Weistroffer, Nicholas McComb, Sindhu Karnam, Adel Azghadi, Kevin S. Murnane, Mark S. Baron, Kathryn L. Holloway

## Abstract

**Background:** Deep brain stimulation (DBS) of the nucleus basalis of Meynert (NBM) has been preliminarily investigated as a potential treatment for dementia. The degeneration of NBM cholinergic neurons is a pathological feature of many forms of dementia. Although stimulation of the NBM has been demonstrated to improve learning, the ideal parameters for NBM stimulation have not been elucidated. This study assesses the differential effects of varying stimulation patterns and duration on learning in a dementia rat model.

**Methods:** 192-IgG-saporin (or vehicle) was injected into the NBM to produce dementia in rats. Next, all rats underwent unilateral implantation of a DBS electrode in the NBM. The experimental groups consisted of i-normal, ii-untreated demented, and iii-demented rats receiving NBM DBS. The stimulation paradigms included testing different modes (tonic and burst) and durations (1-hr, 5-hrs, and 24-hrs/day) over 10 daily sessions. Memory was assessed pre- and post-stimulation using two established learning paradigms: novel object recognition (NOR) and auditory operant chamber learning.

**Results:** Both normal and stimulated rats demonstrated improved performance in NOR and auditory learning as compared to the unstimulated demented group. The burst stimulation groups performed better than the tonic stimulated group. Increasing the daily stimulation duration to 24-hr did not further improve cognitive performance in an auditory recognition task and degraded the results on a NOR task as compared with 5-hr.

**Conclusion:** The present findings suggest that naturalistic NBM burst DBS may offer a potential effective therapy for treating dementia and suggests potential strategies for the reevaluation of current human NBM stimulation paradigms.

## 1. INTRODUCTION

The pathology of Alzheimer’s (AD) and Parkinson’s disease dementia (PDD) are marked by the accumulation of abnormal proteins in the brain (beta amyloid and hyperphosphorylated tau in AD and alpha-synuclein in PDD) as well as cholinergic depletion and progressive brain atrophy (Braak and Braak, 1991; Gratwicke et al., 2013; Jack et al., 2013; Pagonabarraga and Kulisevsky, 2012). A recent metanalysis of anti-amloyid antibody trials, including Lecanemab and Donanemab, have demonstrated an average reduction in CDR-SB rates of 9% to 21% across all studies compared with progression rates on placebo(Andrews et al., 2019; Teipel et al., 2024). Although this is a step in the right direction, “these effects are below the minimal clinically important difference in CDR-SB for mild cognitive impairment (MCI) and mild AD cases, based both on clinician judgement and a statistical criterion of half the baseline’s SD” (Andrews et al., 2019; Teipel et al., 2024). Furthermore, there is no evidence that Lecanemab or Donanemab reverse cognitive deficits. An additional hallmark of AD and PDD is the dramatic and progressive loss of the neurons in nucleus basalis of Meynert (NBM) and with it, the extensive cholinergic supply to the cortex (Candy et al., 1983; Etienne et al., 1986; Gratwicke et al., 2013; Perry et al., 1985). The NBM plays a critical role in learning, memory and other higher-level cognitive processes (Berger-Sweeney et al., 2000; Butt et al., 2003; Chudasama et al., 2004; Linster et al., 2001). The release of neocortical Acetylcholine (ACh) from the NBM critically regulates neocortical excitability, and provide a faciliatory platform for the neuroplastic integration of new sensory inputs within memory engrams (Weinberger et al., 2013, 2006). NBM activity also induces the release of cortical nerve growth factor (NGF) which enriches and thereby sustains the local cholinergic neurons, as well as the cortical regions to which the NBM projects (Bambico et al., 2015; Stone et al., 2011). Thus, the degenerative loss of NBM activity results in a downward spiral of decreased cortical activation and further cholinergic loss. Cholinergic replacement medications are not only limited by providing only modest therapeutic benefits, but they also do not impact this downward pathological spiral. In distinction, electrical stimulation of the NBM has been shown to raise ACh and NGF levels in the NBM and in the affected cortex and to improve cognition in animal models (Berger-Sweeney et al., 2000; Butt et al., 2003; Freund et al., 2009; Kurosawa et al., 1989; Linster et al., 2001; Suga, 2020; Weinberger et al., 2013, 2006). NBM stimulation approaches to enhancing cognitive functioning are potentially complementary to immunotherapeutic targeting of amyloid in the case of AD. For example, NBM stimulation in Amyloid-b precursor protein/Presenilin1 (APP/PS1) transgenic mice has resulted in downregulation soluble Ab40 and Ab42 in the hippocampus (Huang et al., 2019) as well as in chronic entorhinal cortex DBS in 3xTg mice, another well-established animal model of AD (Mann et al., 2018).

The promising experimental results in animals have led several investigators to explore the use of deep brain stimulation (DBS) of NBM for the treatment of dementia in clinical trials (Barnikol et al., 2010; Freund et al., 2009, 2009; Gratwicke et al., 2018; Kuhn et al., 2015a; Lee et al., 2019; Nombela et al., 2019). Similar to the anti-amyloid antibody trials, these studies showed a slowing of the rate of progression compared with controls, but not a dramatic effect. In the Kuhn trial of NBM DBS in 6 AD patients, the group mean worsened by 3 points in the AD assessment score (ADAS-Cog) over the course of the year (Kuhn et al., 2015a) with tonic stimulation. This represents a preliminary improvement over the decrease of 4.5 seen in a large long-term study of progression of AD in patients treated with anti-dementia drugs (Scarmeas et al., 2007), the 4.2 point worsening seen in the Phase 1 fornix DBS trial, and the complete lack of benefit reported in the Phase 2b Fornix trial. It is similar to the 10 point worsening at 76 weeks in the Donanemab trial compared with 13 point worsening in the placebo group(Sims et al., 2023). A trial of tonic NBM DBS in PDD patients yielded 2 of 6 patients with significant improvement in hallucinations but otherwise no significant improvements. The modest, inconsistent benefits of NBM tonic DBS in human trials to date contrast with the clear improvements in learning seen most effectively with NBM burst stimulation in animal dementia models. As we have previously published (Kumbhare et al., 2018), the differences in stimulation parameters between animal and human studies may be a critical factor for the underwhelming responses in the human trials, which have strictly investigated continuous 24-hour tonic stimulation. Burst stimulation is more characteristic of the normal physiological oscillatory patterns of the brain (Akam and Kullmann, 2014; Buzsáki and Wang, 2012; Fountas and Shanahan, 2018; Headley and Paré, 2017). Brief bursts of NBM stimulation in animal models have demonstrated that the induced release of Ach is associated with gamma oscillations and protein synthesis, and this has been associated with learning. Constant stimulation is likely to have effects on sleep, as cholinergic activity plays a key role in sleep transitions (Irmak and de Lecea, 2014). Lastly, NBM activity induces an attentive state, and this may induce stress if maintained constantly. On this background, with a goal to advance more physiological and effective approaches for NBM DBS, we investigated the effects of different modes and durations of NBM stimulation on cognition in a cholinergic toxin model of dementia in rats.

## 2. MATERIALS AND METHODS

### 2.1. Animals

All experiments were approved and monitored by the Institutional Animal Care and Use Committee of the Richmond Veterans Affairs Medical Center and performed in accordance with regulatory guidelines. Long Evans wild-type (LE-WT) rats were used for this study. Animals were initially obtained from Charles River, MA, USA and maintained as an in-house breeding colony in the Richmond Institute for Veteran’s Research animal facility. The rats were housed on a 12-hr light/ 12-hr dark cycle with food and water ad libitum. Animals were housed in groups of 2 or 3 per cage before procedures and were single-housed post-surgery. To prevent additional gender related variability, only male rats, 8-48 weeks old and 200-720 g at the time of stereotactic injection of the tracers were used. All experiments were conducted in strict accordance with the ARRIVE guidelines. All data were reviewed by multiple team members to ensure its validity and to minimize operator biases.

### 2.2. 192-Ig-G saporin model refinement

#### 1. Intraparenchymal injections

To define the amount of 192-IgG saporin to achieve 70-80% destruction of cholinergic population in NBM in our lab, we performed a series of intraparenchymal injections with varying concentrations. Animal surgeries were carried out under 1.5-3% general isoflurane anesthesia (with 1 L/min O_2_) in adult rats. Two burr holes (2 mm) were made on either side of the midline of the skull centered over the NBM (AP −2.3, ML ±3.3 mm; fig. 1A). The location of the NBM targets were confirmed using electrophysiological mapping using a Thomas Recording heptode assembly. Once the trajectory was confirmed, 192-IgG saporin (Advanced Targeting Solution) was bilaterally injected in NBM over 5 min at a depth of 8 mm from the surface of the brain using a Hamilton syringe (65459-02, Neuros 2 μl) and syringe pump (fig. 1A). The following amounts of 192-IgG saporin: 0.0 µg (control), 0.15 µg, 0.3 µg, 0.50 µg, and 0.75 µg in 0.5 µl of Dulbecco’s phosphate buffered saline (dPBS) were injected over 5 min in NBM in both hemispheres in two rats per concentration (n = 10 total rats). A 10 min period was allocated before withdrawing the injection cannula. The incision was sutured, and the rat was returned to its home cage and was regularly monitored for potential health complications. Refer to Table 1 for a summary of rats and procedures used for the study.

**Fig 1.**
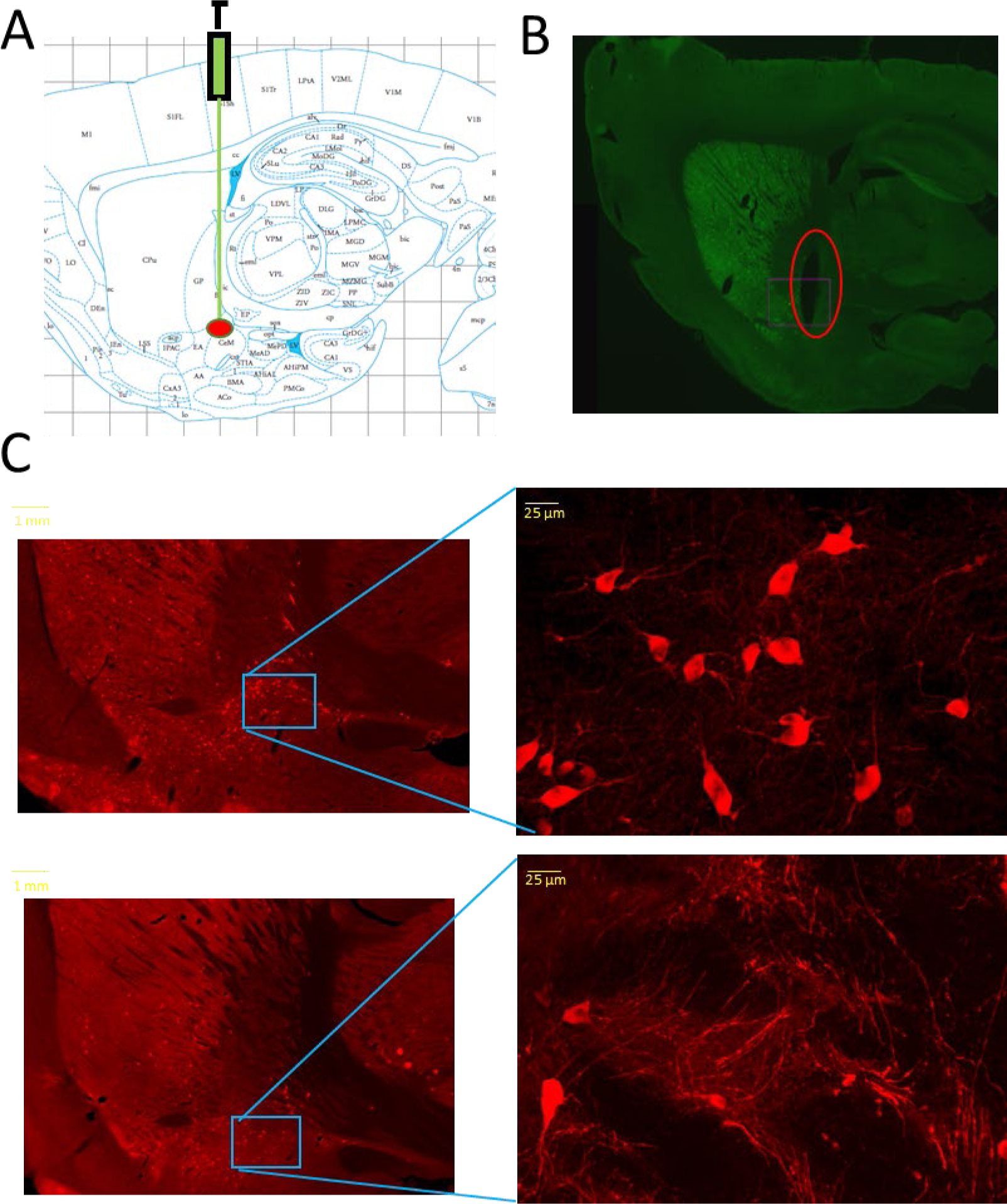
Stereotactic injection of IgG saporin and implantation of stimulation electrode in rats. **(A)** Illustration of Intraparenchymal injection of saporin in NBM using micropipette. The red dot indicates the injection target. **(B)** A representative sagittal image of one of the rat brain showing track damage (red circle) due to implanted DBs electrode. **(C)** Sagittal brain sections stained with choline acetyltransferase (ChAT) labelling the cholinergic cell population, focusing on NBM area. Top panel: Control brain with injection of dPBS. The NBM is populated with cholinergic neurons. Bottom pannel: Saporin injected brain. The NBM area has a significant loss of cholinergic neurons and unhealthy nerve fibers can be seen. The right panels show magnified images of the NBM region.

**TABLE 1.**
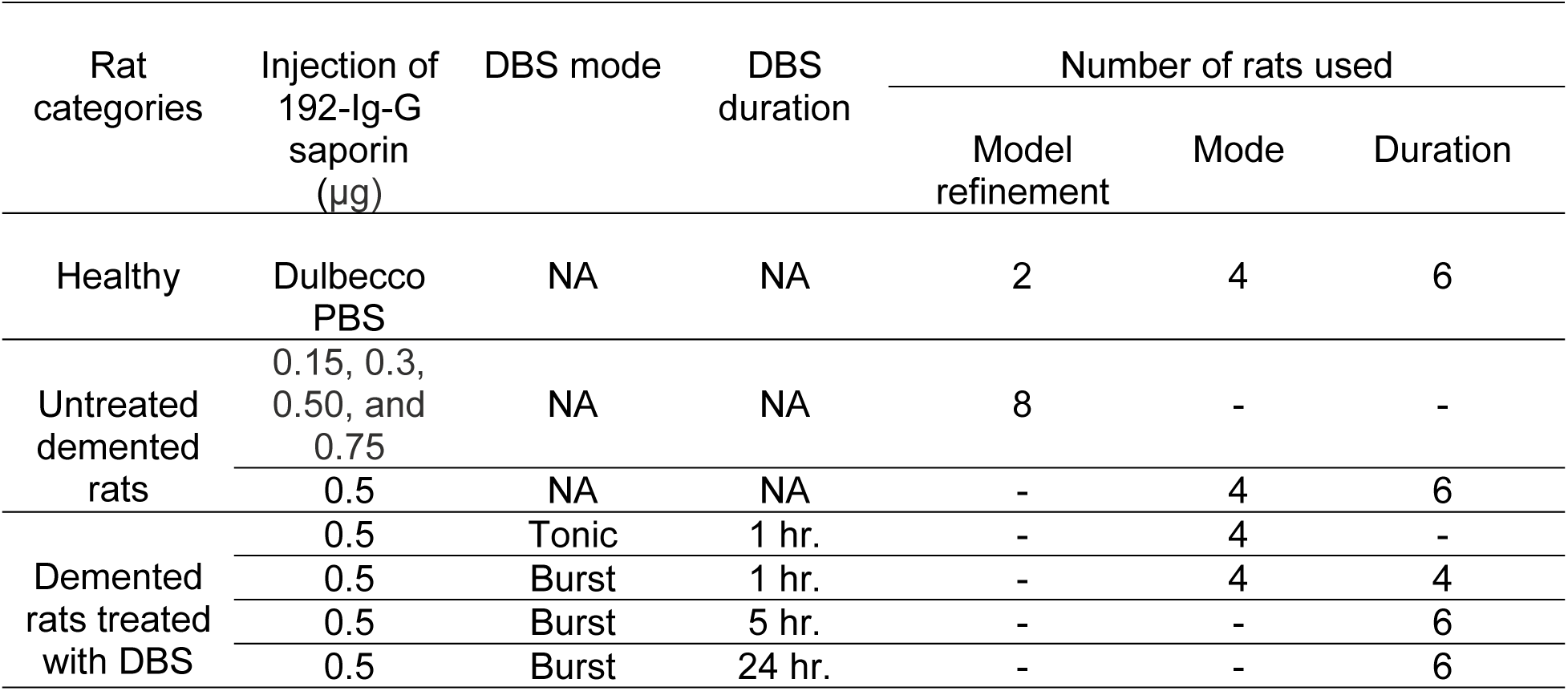
Details of the rats used for model refinement, Comparison of DBS modes and DBS durations.

#### 2. Histology

The rats were euthanized on day 6 of the injection using pentobarbital and perfused with 4% paraformaldehyde (PFA). The brain was harvested and initially stored in PFA overnight and then moved to a 1xPBS and 30% sucrose solution. The right hemisphere of each brain was then set in Tissue-Tek O.C.T. Compound (Sakura Finetek #4583), frozen at −80°C, and sectioned into 50 µm sagittal slices via cryostat (Leica CM1860, Leica Biosystems). The brain sections were stored in a solution of 1xPBS and 0.2% sodium azide until staining was performed. Brain sections were selected for staining based on track damage and laterality of the brain section based on a rat brain atlas, focusing on the NBM area of the brain (Paxinos and Watson, n.d.). A total of 4-5 sagittal brain sections (100 µ) in the region of interest (ROI) defined between L3.2 -L3.7 were stained with antibody choline acetyltransferase (ChAT) to assess the cholinergic population in the NBM (Yi et al., 2015). On day 1, the free-floating brain sections were first washed in 1x PBS for 5 minutes, twice. Then the sections were quenched with 3% hydrogen peroxide for 1 hour, rinsed with a PBS + 0.3% Triton100 (Thermo Scientific #85111) solution for 10 minutes, and then blocked for 2 hours at room temperature in a 2.5% normal donkey serum (abcam ab7475) and 0.3% Triton100(Casini et al., 2018). The brain sections were then incubated for 48 hours at 4 °C in a primary ChAT antibody solution at a concentration of 1:200, diluted with PBS + 0.3% Triton100, 5% normal donkey serum, and the anti-choline acetyltransferase antibody (Sigma-Aldrich AB144P). After primary antibody incubation, the sections were washed with PBS + 0.3% Triton100, and blocked for 2 hours at room temperature with the same normal donkey serum and Triton solution as used previously. Then, the brain sections were incubated for 3 hours at room temperature, or overnight at 4°C, in the light sensitive secondary antibody solution containing PBS + Triton, normal donkey serum, and 568 donkey anti-goat secondary antibody (Alexa Fluor A11057) in a dilution of 1:500. After the secondary antibody incubation, the brain sections were washed 3 times in PBS for 5 minutes. The tissue was then mounted on a labeled slide (Fisherbrand 22-037-246), 4 sections per slide, coverslipped with Prolong Gold antifade reagent (Invitrogen P36934), and the edge was sealed with clear nail polish. The slides were allowed to set for 24 hours before imaging. Careful attention was paid during the secondary antibody phase due to the light sensitive nature of the stain.

#### 3. Imaging

Cholinergic cell density in the NBM for each group was quantified using a BZ-X800 Keyence fluorescence microscope. Images were obtained at 20x to see the density of the cholinergic population, and at 40x to assess the quality of the cells (fig. 1C). Cholinergic neurons in the NBM region were counted at 20x using the anterior commissure as the bottom border and counting up to half of the globus pallidus (GP). The total number of cholinergic cells contained within these borders were manually counted for each brain section.

### 2.3. Rat implantation surgery

Based on the results from testing varying dosing (see section 3.1 below), 192-IgG saporin 0.5 µg in 0.5 µl DPBS per side was injected bilaterally in NBM. The implantation surgery for testing effects of stimulation parameters involves similar intraparenchymal injection of 192-IgG saproin in NBM as described in section 2.2.1 above. Control rats were injected bilaterally with 0.5 µl of pure DPBS. Both stimulation and control groups also underwent stereotactic unilateral DBS electrode (0.203 mm Tungsten electrode, invivo1 Inc.) implantation in the NBM auditory territory (Weinberger et al., 2013, 2009) (coordinates: AP: −2.3, ML +3.3 mm, DV 8 mm). Figure 1B illustrates an example of the electrode track damage in an implanted rat. In addition, six holes were drilled into the skull and stainless-steel micro-screws (Invivo1) were secured for anchoring and to secure a return electrode and a reference. Next, all the electrode connectors were assembled on a socket contact and a pedestal backmount (Invivo1) and were secured with bone cement (Lang Ortho-Jet™ Package). A dust cap was subsequently placed over the socket contact. Prior to discontinuing the anesthesia, bupivacaine (0.1-0.55 ml) was injected along the incision. The rat was then returned to its home cage and was regularly monitored.

#### Post-care

Generally, the SAP injected rats became ill and were weaker compared to the control rats during post-surgery recovery and needed extra care and health monitoring for 4-6 days. Figure 2A indicates the schematics of the experimental timeline following the implantation surgery.

**Fig 2.**
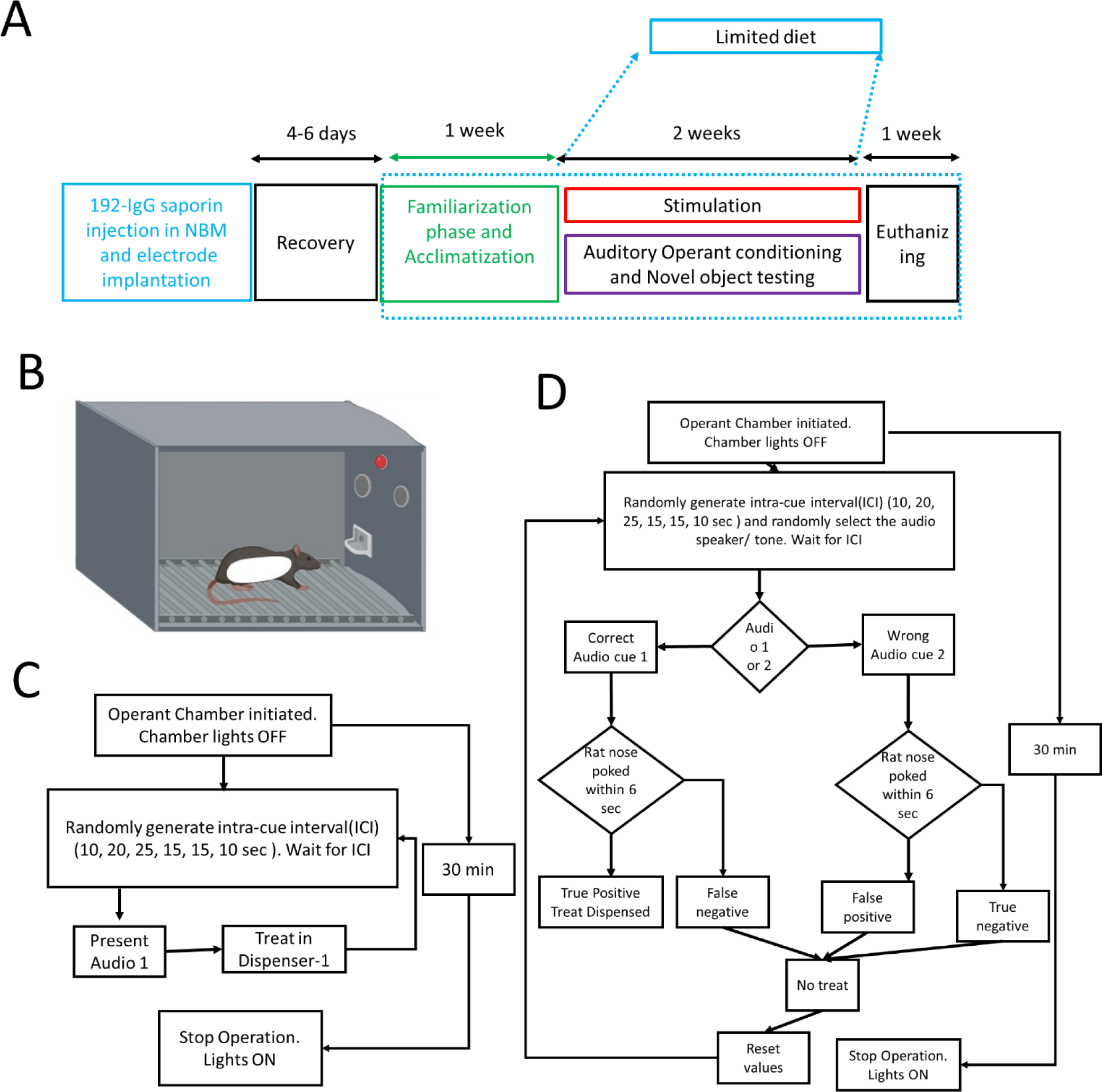
Experimental timeline and operant chamber paradigms. **(A)** Timeline of the stimulation experiments employed in this study. (B) Operant chamber assembly indicating the arrangement of the audio speakers, house light, and the treat dispenser. (C) Flow chart depicting the acclimatization paradigm for auditory training paradigm. (D) Details of the auditory learning training paradigm.

### 2.4. Auditory cue based training

#### Food restriction

After 4-6 days to recover from the effects of injections, to increase motivation for the food reward task, all rats were placed on food restriction until they reached 85–90% free-fed body weight. Weights were closely monitored while recovering from surgery and assured to be stable prior to beginning food restriction. The animals were weighed daily and the food was weighed and distributed accordingly. Water was given ad libitum.

#### Operant chamber

A Med-Associates (St. Albans, VT) operant chamber (30 X 30 X 35 cm^3^) was used for an auditory behavioral paradigm (fig 2B). The chamber contains (i) a 3 cm poking holes attached with a treat dispenser with sensors for poke detection; (ii) two mounted speakers used to deliver conditioned stimulus tones ((i) frequency = 40 KHz and intensity = 80 dB SPL; and *(ii) frequency = 20 KHz; intensity = 80 dB SPL)) and an incandescent light.

#### Acclimatization

During the first week of limited diet, the rats were acclimatized to the auditory behavioral testing environment. The first three sessions (1 session/ day) include placing the rat in the operant chamber for 15 min with 10 treats already present in each of the dispensers. The next three sessions (30 min/ session/ day) include a pre-training paradigm (fig 2C). This includes acclimatizing the rat to a paradigm where it is presented with an audio tone (40 KHz) followed by a treat in the dispenser. 2 rats that didn’t show motivation to finish the dispensed treats were removed from the study.

#### Auditory training

During the auditory paradigm (fig. 2D) the rats were trained to poke their nose in a hole (40 Hz) when the familiar tone is heard. The rats were tested (30 min/session, 5 days a week) on the audio-cue based learning paradigm for two weeks (fig 2D). Two audio tones (40 KHz (rewarded) and 20 KHz (non-rewarded)) were randomly presented at random intervals (10, 15, or 25 sec). The rats were rewarded with a food pellet for nose poking in the treat dispenser within 6 sec of the 40 KHz cue. Failing to respond within 6 secs of the reward cue, responding to the incorrect (20 Khz) auditory cue, and indiscriminate nose poking all equally lowered the accuracy score.

### 2.5. Electrical Stimulation of NBM

In conjunction with the auditory training, the NBM was directly stimulated, beginning 2-3 hrs after each behavioral training session. While in the animal’s home cage, the head implant connector (invivo1) was connected via a commutator (6-channel commutator, In vivo) to an isolated stimulator (AM systems, MODEL 4100). For two weeks, 5 days per week, either continuous 50 Hz tonic (pulse duration = 200 µs) or burst stimulation (50 pulses/burst, pulse duration = 200 µs, intra-burst freq. = 100 Hz) was delivered for 1 hr, 5 hrs, or 24 hrs per session. Cathodic monopolar square wave pulses (50% duty cycle) were delivered through the implanted Invivo1 tungsten electrodes at an amplitude of 80-100µA. For rats receiving 24 hr stimulation, the implant was briefly disconnected from stimulation during the behavioral testing. Control rats were connected to the commutator, but no stimulation was delivered. In-between stimulation sessions, the connector was disconnected and the dust cap was placed. Five additional rats were euthanized after their cranial implant loosened or detached before the completion of training sessions and were excluded from the study.

### 2.6. Novel object recognition (NOR) testing

Novel object recognition (NOR) is the benchmark test of recognition memory in rodents (Antunes and Biala, 2012). The test takes place in an open field (104 square inch area). A video camera (Basler) was placed above the open field to videotape each trial. NOR testing began within 2-3 days of completing the 2 weeks of NBM stimulation (and after completing the auditory learning testing) and was completed over two consecutive days. The day prior to initiating NOR testing, to reduce neophobic responses, the rats were habituated to the empty open field for 30 min. During the initial daily familiarization phase, a rat was placed into the field for 5 min and allowed to explore and become familiar with two identical objects placed in the chamber approximately 38 cm from opposite corners of the open field. The animal was then removed, and the objects and the open field were wiped down with 70% alcohol solution to avoid scent marking. Next, 4 hrs after the familiarization phase, a testing phase began in which one of the familiar objects was replaced by a novel object and the animal was again allowed to explore the open field for 5 min (Antunes and Biala, 2012; Aubele et al., 2008; Frumberg et al., 2007). Offline, video analysis software (Noldus) was used to assess the time spent exploring each of the objects in each phase. A discrimination index (DI) was calculated to assess novel object recognition as the outcome measure (Antunes and Biala, 2012), as described in section 2.8.

### 2.7. Histology and Imaging

One week after completing stimulation and control investigations, the rats were euthanized as described above (under section 2.2). The rats’ brains were processed and imaged as also described above and the intended 192-IgG saporin-induced tissue damage was verified.

### 2.8. Data Analysis

#### 1. Performance metric for auditory paradigm

For the auditory paradigm, performance was determined using accuracy and precision metrics based on components of confusion matrix (true positive, false positive, true negative, and false negative). Accuracy was calculated using the following formula:

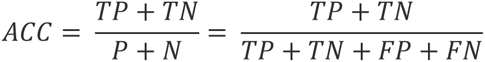

A true positive (TP) is the number of times the rat promptly (within 6 sec) poked their nose in the dispenser after the audio tone. A false negative (FN) is the number of times the audio cue was presented but the rat failed to respond within the time limit. f the rats poked their nose in the dispenser prior to the audio tone, it was counted as a false positive (FP). If the rat did not respond at all to the tone, this was counted as a true negative (TN).

#### 2. Performance metrics for NOR paradigm

A discrimination index (DI) was used to assess object recognition. The DI compared the time spent with a novel versus with a familial object as follows:

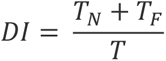

where T_N_ is time spent exploring the novel object, T_F_ is time spent exploring the familiar object, and T is total time spent in the field. Demented rats will treat the 2 objects similarly approaching the null value, whereas non-demented rats will spend more time with the new object departing from the null value.

### 2.9. Statistical analysis

MATLAB was used for statistical analyses. Comparisons of performances between multiple groups during auditory and NOR testing were assessed using Kruskal-Wallis test followed by Tukey’s Honestly significant difference (HSD) multiple comparison test. A probability value of < 0.05 was considered statistically significant for comparisons between groups.

## 3. RESULTS

Refer to Table 1 for details of all the rats used in this study.

### 3.1. Development of 192-IgG saporin model in our lab

Initially, as described above, 10 rats were injected with 192-IgG saporin in the NBM bilaterally with following concentrations: 0.0 µg (control), 0.15 µg, 0.3 µg, 0.50 µg, and 0.75 µg dissolved in 0.5 µl of Dulbecco’s PBS (n = 2 rats per concentration). The ChAT count revealed a total loss of 55 ± 10%, 67 ± 8%, 80 ± 12%, and 100%, respectively of cholinergic population in the injected region, with increasing concentrations of saporin compared to the controls (mean of 39.6 ChAT+ cells/ 100µ thick slice; fig. 1C). Therefore, to assure extensive, but not complete depletion of cholinergic neurons in the NBM, we chose to inject 0.5 µg of saporin for our subsequent behavioral studies.

### 3.2. Effect of DBS mode on learning

#### Experimental groups

To compare the effects of DBS mode (burst vs tonic), four experimental groups (n= 4 rats per group) were assessed: demented rats with i) no stimulation, ii/iii) 50 Hz tonic and delta-gamma burst stimulation, 1 hr/ day, 5 days/ week for 2 weeks and iv) dPBS injected normal control rats.

#### Auditory operant chamber learning

During the auditory training period, healthy control rats (n= 4) showed a gradual improvement in the ability to correlate an audio tone with treat dispensation over the course of training sessions. In contrast, demented rats as a group showed no change or reduced performance over the sessions. A Kruskal-Wallis test revealed statistically significant difference in extent of improvement in accuracy between at least two of the four groups (F (3,12) = 5.78, p =0.011). Both burst and tonic stimulated demented rats showed improvement from baseline in audio task performance over time (fig 3A). Tukey’s HSD Test (fig 3C) for multiple comparisons supported that the improvement was significantly different between normal and dementia rats (p = 0.0304, 95% C.I. = [2.0 43.62]). The extent of the improvement seen with burst DBS was statistically significant with respect to that of the demented rats (p < 0.0201, 95% C.I. = [3.66 45.27]). However, there was no statistically significant difference in extent of improvement between tonic DBS and non-stimulated demented rats (p=0.722).

**Fig. 3.**
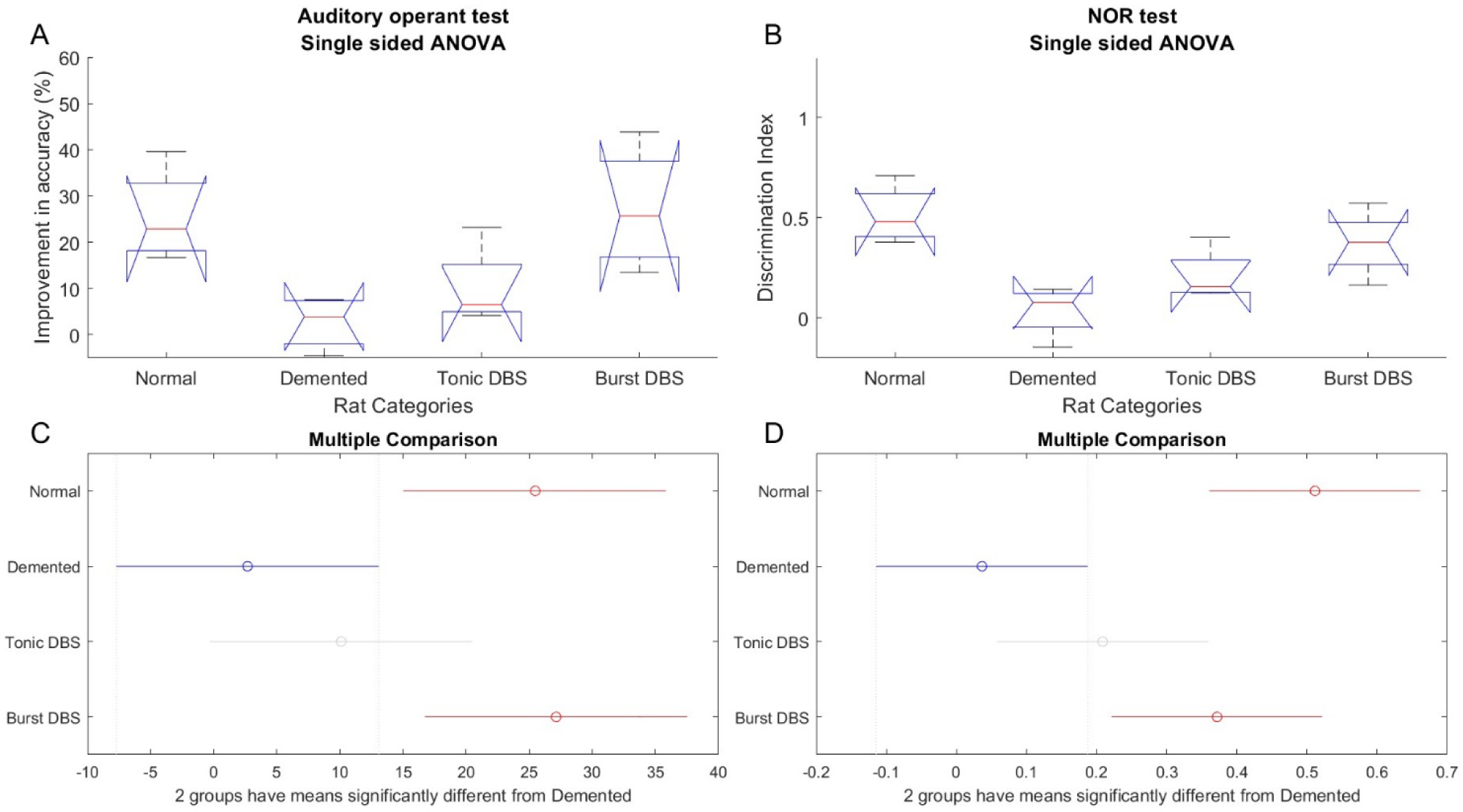
Differential improvement in auditory learning and NOR induced by different stimulation modes. (a) Box and whisker plot showing Kruskal-Wallis test of different categories of rats showing improvement in accuracy from first till last auditory training session. (B) Box and whisker plot showing Kruskal-Wallis test of different categories of rats showing discrimination index for novel object recognition test evaluated after the 2 weeks stimulation session. On each box, the central mark is the median and the edges of the box are the 25th and 75th percentiles. The whiskers extend to the most extreme data points that are not considered outliers. The outliers are plotted individually using the ‘+’ symbol. (C) and (D) are results of Tukey’s Honestly significant difference (HSD) for data from (A) and (B) respectively.

#### Novel object recognition

A Kruskal-Wallis test revealed significant differences in DI for NOR between at least two of the four groups (F (3,12) = 8.17, p =0.0031). Non-stimulated demented rats showed evidence of reduced general memory (fig 3C) with a DI of 0.075, which is near to the null reference (DI = 0; fig 3C). The demented rats performed significantly worse than healthy rats (p = 0.0281, 95% C.I. = [−0.6360 −0.0337]). The DI values for both tonic and burst stimulated rats trended towards that of normal rats, however only the burst DBS rats performed significantly better than the unstimulated demented group (p = 0.0281, 95% C.I. = [−0.6360 - 0.0337]). Although DI scores were lower in the burst compared to tonic stimulated, the differences were not statistically significant (p = 0.12). Refer to Table 2 for detailed HSD multiple comparison test results.

**Table 2.** Results of HSD multiple comparison test for DBS-mode comparison.

| Test | Group | Compared group | Lower limit | Difference | Upper limit | p-value |
| --- | --- | --- | --- | --- | --- | --- |
| Auditory operant test | Normal | Demented | 2 | 22.81 | 43.62 | **0.03** |
|  | Normal | Tonic stimulated | -5.42 | 15.39 | 36.2 | 0.18 |
|  | Normal | Burst stimulated | -22.47 | -1.66 | 19.15 | 0.99 |
|  | Demented | Tonic stimulated | -28.22 | -7.42 | 13.39 | 0.72 |
|  | Demented | Burst stimulated | -45.27 | -24.47 | -3.66 | **0.02** |
|  | Tonic stimulated | Burst stimulated | -37.86 | -17.05 | 3.76 | 0.12 |
| NOR | Normal | Demented | 0.17 | 0.48 | 0.78 | **0.001** |
|  | Normal | Tonic stimulated | 0 | 0.3 | 0.6 | **0.05** |
|  | Normal | Burst stimulated | -0.16 | 0.14 | 0.44 | 0.54 |
|  | Demented | Tonic stimulated | -0.47 | -0.17 | 0.13 | 0.37 |
|  | Demented | Burst stimulated | -0.64 | -0.34 | -0.03 | **0.03** |
|  | Tonic stimulated | Burst stimulated | -0.46 | -0.16 | 0.14 | 0.41 |

### 3.3. Effect of DBS duration on learning

#### Experimental groups

Since burst, but not tonic stimulation, produced significant improvement in auditory behavioral performance, burst stimulation was utilized to investigate the effects of stimulation duration. These studies comprised of five experimental groups: i) unstimulated demented (n = 6 rats), ii/iii/iv) 1 hr/ (n = 4, from section 3.4), 5 hrs/ (n = 6), and 24 hrs/ day stimulation (n = 6), and v) normal controls (n= 6).

#### Auditory operant chamber learning

A Kruskal-Wallis test revealed significant differences in accuracy improvement between at least two of the five groups (F (4,25) = 5.45, p <0.003). Tukey’s HSD Test for multiple comparisons established that unstimulated demented rats experienced deficient learning compared to normal rats (8.0% vs 20.8% improvement, p = 0.0304, 95% C.I. = [0.68 24.84]); Fig 4A and C). Burst stimulated rats, irrespective of the duration of daily stimulation demonstrated greater improvement in learning accuracy over time compared to that of unstimulated demented rats (1-hr (n = 4, median 31.5 %), 5 hr burst (n = 6, 27.8 %), and 24-hr burst (n=6, 35.7 % improvement). These differences reached significance for all three stimulated rat categories (Tukey’s HSD Test for multiple comparison, all p< 0.05. Normal rats learned to respond to the tone more quickly than demented rats (% improvement in response time of normal rats was 47.59%, vs demented rat was 8.39%). The response time improvement in all of the stimulated rats was similar and was significantly quicker than the demented rats (% improvement ranged from 25.18 to 32.9%; all p < 0.05) but still significantly worse than the normal rats (47.59%; all p < 0.05). Rats stimulated for 5 hr/ day showed the maximum improvement in response time.

**Fig. 4.**
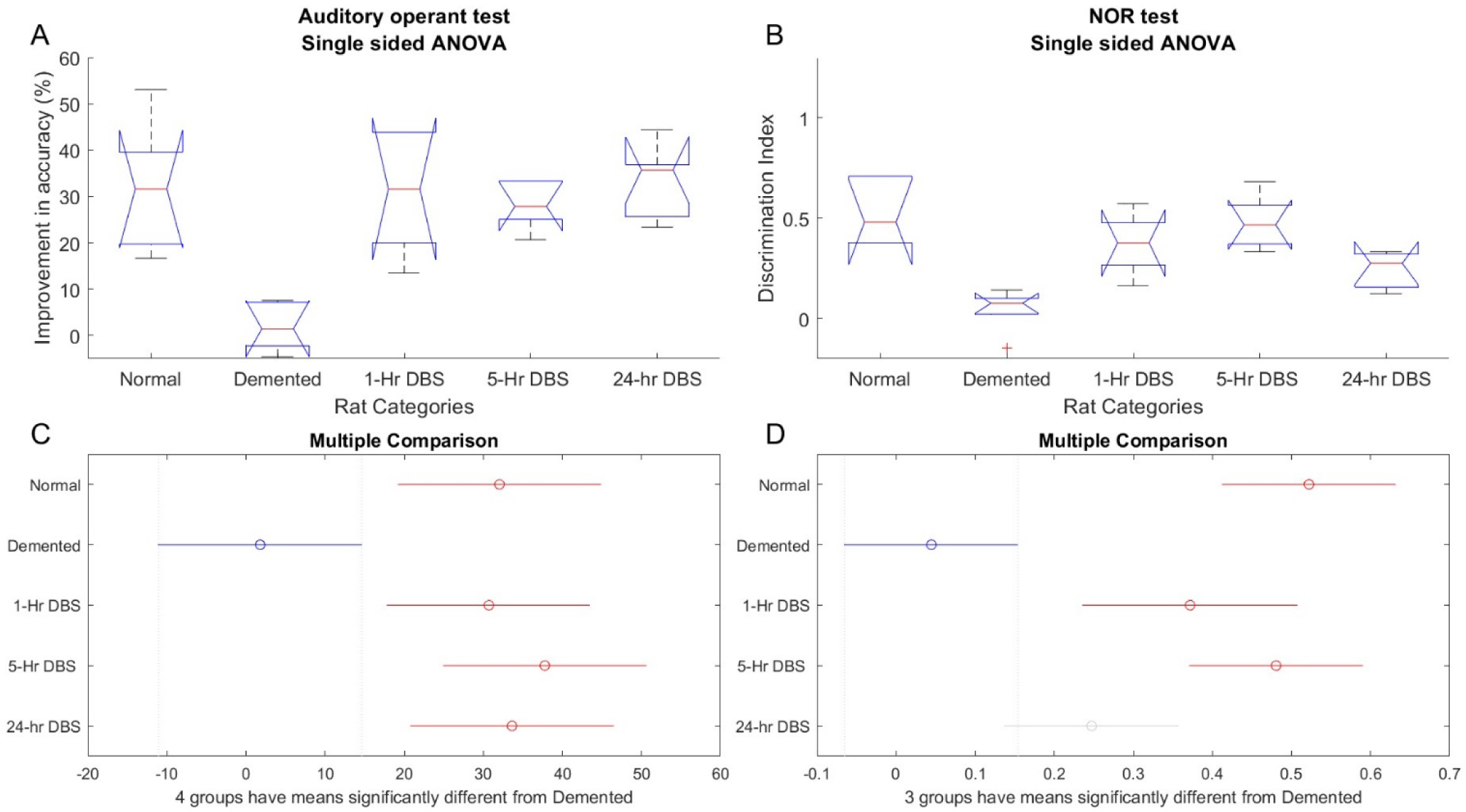
Differential improvement in auditory learning induced by different stimulation duration. (a) Box and whisker plot showing Kruskal-Wallis test of different categories of rats showing improvement in accuracy from first till last auditory training session. (B) Box and whisker plot showing Kruskal-Wallis test of different categories of rats showing discrimination index for novel object recognition test evaluated after the 2 weeks stimulation session. On each box, the central mark is the median and the edges of the box are the 25th and 75th percentiles. The whiskers extend to the most extreme data points that are not considered outliers. The outliers are plotted individually using the ‘+’ symbol. (C) and (D) are results of Tukey’s Honestly significant difference (HSD) for data from (A) and (B) respectively.

#### Novel object recognition

There was a significant difference in DI between at least two of the five groups (F (4,23) = 13.46, p <0.05; fig. 4B). The normal, 1-hr, and 5-hr burst stimulated rats performed significantly better than the demented rats (normal: p <0.05, 95% C.I. = [0.26 to 0.7]; 1-hr DBS: p = 0.01, 95% C.I. = [0.08 0.57]; 5-hr DBS: p < 0.05, 95% C.I. = [0.22 to 0.66]; Fig 4b,d). Although, the 24hr stimulated rats appeared to perform better than demented rats, there was a significantly better performance on the NOR task in the 5-hr. group than with 24-hr. stimulation. This represented the only significant difference between the duration groups. Detailed results of HSD multiple comparison test are shown in table 3.

**Table 3.** Results of HSD multiple comparison test for DBS-duration comparison.

| **Test** | **Group** | **Compared group** | **Lower limit** | **Difference** | **Upper limit** | **p-value** |
| --- | --- | --- | --- | --- | --- | --- |
| Auditory operant test | Normal | Demented | 4.56 | 30.26 | 55.96 | **0.02** |
|  | Normal | 1-hr DBS | -24.37 | 1.34 | 27.04 | 1 |
|  | Normal | 5-hr DBS | -31.43 | -5.73 | 19.98 | 0.96 |
|  | Normal | 24-hr DBS | -27.28 | -1.58 | 24.12 | 1 |
|  | Demented | 1-hr DBS | -54.62 | -28.92 | -3.22 | **0.02** |
|  | Demented | 5-hr DBS | -61.69 | -35.99 | -10.28 | **<0.01** |
|  | Demented | 24-hr DBS | -57.54 | -31.84 | -6.14 | **0.01** |
|  | 1-hr DBS | 5-hr DBS | -32.77 | -7.06 | 18.64 | 0.93 |
|  | 1-hr DBS | 24-hr DBS | -28.62 | -2.92 | 22.79 | 1 |
|  | 5-hr DBS | 24-hr DBS | -21.56 | 4.15 | 29.85 | 0.99 |
| NOR | Normal | Demented | 0.26 | 0.48 | 0.7 | **0.01** |
|  | Normal | 1-hr DBS | -0.1 | 0.15 | 0.4 | 0.39 |
|  | Normal | 5-hr DBS | -0.18 | 0.04 | 0.26 | 0.98 |
|  | Normal | 24-hr DBS | -0.05 | 0.27 | 0.49 | **0.01** |
|  | Demented | 1-hr DBS | -0.57 | -0.33 | -0.08 | **0.01** |
|  | Demented | 5-hr DBS | -0.66 | -0.44 | -0.22 | **0.01** |
|  | Demented | 24-hr DBS | -0.42 | -0.2 | 0.02 | 0.08 |
|  | 1-hr DBS | 5-hr DBS | -0.35 | -0.11 | 0.14 | 0.69 |
|  | 1-hr DBS | 24-hr DBS | -0.12 | 0.12 | 0.37 | 0.57 |
|  | 5-hr DBS | 24-hr DBS | -0.01 | 0.23 | 0.45 | **0.03** |

## 4. DISCUSSION

NBM stimulation primarily affects the neocortical environment by modulating the discharge properties of the target nuclei and by release of ACh and NGF to alter the neocortical excitability. The optimal stimulation paradigm should provide a facilitatory platform for neuroplasticity, aiding memory consolidation(Weinberger et al., 2013, 2006). Our results demonstrate the expected effect of intraparenchymal SAP injection on learning, with demented rats demonstrating significantly lower learning ability and memory in the auditory paradigm and NOR as compared to normal controls; thus, demonstrating that the chosen model is an effective dementia model. NBM stimulation resulted in an improvement in learning in an auditory paradigm that was significantly better than the demented controls. The naturalistic burst mode was relatively more effective than the tonic mode in improving the learning deficits. Learning improvement became more consistent by increasing the duration of NBM stimulation from 1 hour to 5 hours. The 5-hr group, in fact, demonstrated more improvement in both the behavioral (auditory operant as well as NOR) tests from baseline than the pathological rats whereas the 1 hr group only demonstrated significant improvement in NOR. Increasing the stimulation duration to 24 hrs/ day did not cause any additional behavioral improvement.

There are important differences in the stimulation parameters used in all the prior animal work as compared to the human trials. The pilot trials of NBM stimulation in AD, PDD, and Lewy body dementia all employed a tonic stimulation (Freund et al., 2009; Gratwicke et al., 2018; Hardenacke et al., 2016; Kuhn et al., 2015a, 2015a; Maltête et al., 2020; Nombela et al., 2019). This was initially chosen based on the tonic portion of the output pattern of the NBM and has demonstrated some mild benefit in the study patients. Unal et al. found dichotomous cholinergic neuronal population in NBM in rats, which includes highly excitable fast responding neurons and less excitable slow responding tonic neurons (Unal et al., 2012). While the former employs spike frequency adaptation, the latter fires in a consistent slow tonic mode. Although both of these modes are reported to be critical for information processing, Sarter et al. argues that the adaptive phasic burst mode is more associated with learning (Sarter et al., 2009). Our results provide some preliminary support for this theory. There is a growing body of DBS literature suggesting that certain types of burst or irregular stimulation may not only be more efficient, but they may also lead to long lasting effects within the neural circuit (Adamchic et al., 2014; Brocker et al., 2017; Ebert et al., 2014). However, Dudar et al (Dudar, 1975) reported that 50 Hz stimulation was as effective as burst stimulation for the release of ACH. Thus, it is unclear whether these differences are critical in the human application or whether both modes are equally effective. Our results provide preliminary evidence for the effectiveness of burst mode over the tonic.

Most of the previous animal NBM studies have been conducted with short duration trains of stimulation, whereas the prior clinical trials (Baldermann et al., 2018; Barnikol et al., 2010; Freund et al., 2009; Gratwicke et al., 2020, 2018; Hardenacke et al., 2016; Kuhn et al., 2015b; Maltête et al., 2020; Nombela et al., 2019, 2019) were applied to the patients for 24 hours per day 7 days a week. It is not clear whether the constant activation of the NBM is counterproductive, but it is hypothesized that it may introduces potential problems related to the NBM’s significant role in regulating arousal, attention, and sleep cycles (Berger-Sweeney et al., 2000; Butt et al., 2003; Chudasama et al., 2004; Linster et al., 2001). Activation of the NBM results in an attentive state, and although brief periods of attention are beneficial, the effect of continual attention or vigilance is associated with stress (Galinsky et al., 1993; Warm et al., 2008). Sleep is also critical for learning, innate neurogenesis, and may be involved in the clearance of abnormal proteins such as amyloid (García-García et al., 2011; Guzman-Marin et al., 2007; Hairston et al., 2005; Ju et al., 2013; Junek et al., 2010; Kang et al., 2009; Mirescu et al., 2006; Roman et al., 2005; Rothman et al., 2013; Sportiche et al., 2010). Depending on when activation occurs, the NBM is involved in the transition between sleep states to rapid eye movement (REM) sleep or wakefulness (Han et al., 2014; Irmak and de Lecea, 2014; Kalinchuk et al., 2008; Zant et al., 2016). In the absence of a paired stimulus, activation of NBM results in amplification of lesser stimuli(Weinberger et al., 2006). It is unclear whether this will serve to enhance or overload the cognitive network. These differences may be critical to the efficacy of stimulation in humans. Our experiments indicate that the efficacy of DBS improvements is comparable in rats that were stimulated for 24 hrs and that received 5-hr DBS. Thus, increasing duration to 24 hours did not lead to consistent further improvement. This is highly relevant as the clinical trials of NBM DBS have all been conducted with 24 hrs stimulation. Further studies are required in order to understand these potential effects through the comparison of continuous stimulation to stimulation limited to the awake periods.

Further investigation of stimulation parameters is necessary to achieve efficacious sets of NBM-DBS. Many human studies used a low 20 Hz frequency for stimulation of NBM for dementia treatment. Rasmussen suggests that continuous stimulation at 20-50 Hz resulted in maximal ACh release (5). On the other hand, with a train of bursts stimulation, maximal response was observed with intra burst frequency of 100 Hz (36), while 50 Hz intra-burst stimulation halved the response. Liu et al (Liu et al., 2022) found that low frequency (20 Hz) intermittent stimulation (20s on, 40s off) improved learning deficits in rat model of scopolamine (Liu et al., 2022). This paradigm is based on that of a prior work out of Birmingham (Liu et al) that evaluated tonic stimulation in 3 rhesus monkeys during a working memory task and found that their performance decreased with tonic stimulation but improved with an intermittent stimulation pattern of 60 Hz for 20 secs followed by 40 secs of rest. The improvement in the short term was modest, however they found a very interesting progressive improvement in performance over time, that was unexplained. In ongoing work, we are evaluating the long-term effects of stimulation using our current experimental paradigm as well as short-term comparisons of the efficacy of naturalistic burst and 20/40 second intermittent stimulation patterns. Additionally, the histologic effects of stimulation and its relationship to cognitive improvement will need to be evaluated in future experiments. Furthermore, these will need to be more comprehensive studies involving both genders to reveal any potential gender-based variability in the results.

## ACKNOWLEDGEMENTS

We thank Ketan Verma, Jan Hachmann, Satvika Nimmagadda, and Greyson Jadwin for assisting during implantation surgeries and behavioral training. We thank Dr. Laxmikant Deshpande for his expert suggestions on designing the behavioral tasks.

## DATA AND CODE ACCESSIBILITY

The digitized data, and the MATLAB custom scripts will be provided upon request. Correspondence and requests for materials should be addressed to DK.

## CRediT authorship contribution statement

<u>Deepak Kumbhare</u>: Conceptualization, designing behavioral and stimulation paradigms, training staff, performing experiments, Data curation, Data analysis and MATLAB coding, Funding acquisition, Writing – original draft, Writing – review & editing. <u>Megan Rajagopal:</u> Funding acquisition, performing experiments, data collection, Writing – review & editing. <u>Jamie Toms,</u> <u>and Anne Freelin</u>: training staff, performing experiments, data collection, histology and imaging, Writing – review & editing. <u>George Weistroffer</u>: performing experiments, data collection, histology and imaging, technical troubleshooting. <u>Nicholas McComb, Sindhu Karnam, and Adel</u> <u>Azghadi</u>: performing experiments, data collection, histology and imaging, Writing – review & editing. <u>Kevin S. Murnane</u>: Critical review and paradigm design. <u>Mark Baron</u>: Critical review and laboratory support. <u>Kathryn Holloway</u>: Conceptualization, Funding acquisition, Project administration, Supervision, Writing – review & editing.

## Conflict of Interest Statement

The authors declare no competing financial interests.

**No disclaimers**

## REFERENCES

Adamchic, I., Hauptmann, C., Barnikol, U.B., Pawelczyk, N., Popovych, O., Barnikol, T.T., Silchenko, A., Volkmann, J., Deuschl, G., Meissner, W.G., Maarouf, M., Sturm, V., Freund, H.-J., Tass, P.A., 2014. Coordinated reset neuromodulation for Parkinson’s disease: Proof-of-concept study. Mov. Disord. 29, 1679–1684. 10.1002/mds.25923

Akam, T., Kullmann, D.M., 2014. Oscillatory multiplexing of population codes for selective communication in the mammalian brain. Nat. Rev. Neurosci. 15, 111–122. 10.1038/nrn3668

Andrews, J.S., Desai, U., Kirson, N.Y., Zichlin, M.L., Ball, D.E., Matthews, B.R., 2019. Disease severity and minimal clinically important differences in clinical outcome assessments for Alzheimer’s disease clinical trials. Alzheimers Dement. Transl. Res. Clin. Interv. 5, 354–363. 10.1016/j.trci.2019.06.005

Antunes, M., Biala, G., 2012. The novel object recognition memory: neurobiology, test procedure, and its modifications. Cogn. Process. 13, 93–110. 10.1007/s10339-011-0430-z

Aubele, T., Kaufman, R., Montalmant, F., Kritzer, M.F., 2008. Effects of gonadectomy and hormone replacement on a spontaneous novel object recognition task in adult male rats. Horm. Behav. 54, 244–252. 10.1016/j.yhbeh.2008.04.001

Baldermann, J.C., Hardenacke, K., Hu, X., Köster, P., Horn, A., Freund, H.-J., Zilles, K., Sturm, V., Visser-Vandewalle, V., Jessen, F., Maintz, D., Kuhn, J., 2018. Neuroanatomical Characteristics Associated With Response to Deep Brain Stimulation of the Nucleus Basalis of Meynert for Alzheimer’s Disease: DBS FOR ALZHEIMER’S DISEASE: ANATOMICAL PREDICTORS. Neuromodulation Technol. Neural Interface 21, 184–190. 10.1111/ner.12626

Bambico, F.R., Bregman, T., Diwan, M., Li, J., Darvish-Ghane, S., Li, Z., Laver, B., Amorim, B.O., Covolan, L., Nobrega, J.N., Hamani, C., 2015. Neuroplasticity-dependent and -independent mechanisms of chronic deep brain stimulation in stressed rats. Transl. Psychiatry 5, e674. 10.1038/tp.2015.166

Barnikol, T.T., Pawelczyk, N.B.A., Barnikol, U.B., Kuhn, J., Lenartz, D., Sturm, V., Tass, P.A., Freund, H.-J., 2010. Changes in apraxia after deep brain stimulation of the nucleus basalis Meynert in a patient with Parkinson dementia syndrome. Mov. Disord. Off. J. Mov. Disord. Soc. 25, 1519–1520. 10.1002/mds.23141

Berger-Sweeney, J., Stearns, N.A., Frick, K.M., Beard, B., Baxter, M.G., 2000. Cholinergic basal forebrain is critical for social transmission of food preferences. Hippocampus 10, 729–738. 10.1002/1098-1063(2000)10:6<729::AID-HIPO1010>3.0.CO;2-M

Braak, H., Braak, E., 1991. Neuropathological stageing of Alzheimer-related changes. Acta Neuropathol. (Berl.) 82, 239–259. 10.1007/BF00308809

Brocker, D.T., Swan, B.D., So, R.Q., Turner, D.A., Gross, R.E., Grill, W.M., 2017. Optimized temporal pattern of brain stimulation designed by computational evolution. Sci. Transl. Med. 9, eaah3532. 10.1126/scitranslmed.aah3532

Butt, A.E., Schultz, J.A., Arnold, L.L., Garman, E.E., George, C.L., Garraghty, P.E., 2003. Lesions of the rat nucleus basalis magnocellularis disrupt appetitive-to-aversive transfer learning. Integr. Physiol. Behav. Sci. Off. J. Pavlov. Soc. 38, 253–271. 10.1007/BF02688857

Buzsáki, G., Wang, X.-J., 2012. Mechanisms of Gamma Oscillations. Annu. Rev. Neurosci. 35, 203–225. 10.1146/annurev-neuro-062111-150444

Candy, J.M., Perry, R.H., Perry, E.K., Irving, D., Blessed, G., Fairbairn, A.F., Tomlinson, B.E., 1983. Pathological changes in the nucleus of Meynert in Alzheimer’s and Parkinson’s diseases. J. Neurol. Sci. 59, 277–289. 10.1016/0022-510x(83)90045-x

Casini, A., Vaccaro, R., Toni, M., Cioni, C., 2018. Distribution of choline acetyltransferase (ChAT) immunoreactivity in the brain of the teleost Cyprinus carpio. Eur. J. Histochem. EJH 62. 10.4081/ejh.2018.2932

Chudasama, Y., Dalley, J.W., Nathwani, Falguni, Bouger, P., Robbins, T.W., Nathwani, Falgyni, 2004. Cholinergic modulation of visual attention and working memory: dissociable effects of basal forebrain 192-IgG-saporin lesions and intraprefrontal infusions of scopolamine. Learn. Mem. Cold Spring Harb. N 11, 78–86. 10.1101/lm.70904

Dudar, J.D., 1975. The effect of septal nuclei stimulation on the release of acetylcholine from the rabbit hippocampus. Brain Res. 83, 123–133. 10.1016/0006-8993(75)90863-X

Ebert, M., Hauptmann, C., Tass, P.A., 2014. Coordinated reset stimulation in a large-scale model of the STN-GPe circuit. Front. Comput. Neurosci. 8.

Etienne, P., Robitaille, Y., Wood, P., Gauthier, S., Nair, N.P., Quirion, R., 1986. Nucleus basalis neuronal loss, neuritic plaques and choline acetyltransferase activity in advanced Alzheimer’s disease. Neuroscience 19, 1279–1291. 10.1016/0306-4522(86)90142-9

Fountas, Z., Shanahan, M., 2018. Correction: The role of cortical oscillations in a spiking neural network model of the basal ganglia. PLoS ONE 13, e0205472. 10.1371/journal.pone.0205472

Freund, H.-J., Kuhn, J., Lenartz, D., Mai, J.K., Schnell, T., Klosterkoetter, J., Sturm, V., 2009. Cognitive functions in a patient with Parkinson-dementia syndrome undergoing deep brain stimulation. Arch. Neurol. 66, 781–785. 10.1001/archneurol.2009.102

Frumberg, D.B., Fernando, M.S., Lee, D.E., Biegon, A., Schiffer, W.K., 2007. Metabolic and behavioral deficits following a routine surgical procedure in rats. Brain Res. 1144, 209–218. 10.1016/j.brainres.2007.01.134

Galinsky, T.L., Rosa, R.R., Warm, J.S., Dember, W.N., 1993. Psychophysical Determinants of Stress in Sustained Attention. Hum. Factors 35, 603–614. 10.1177/001872089303500402

García-García, F., De la Herrán-Arita, A.K., Juárez-Aguilar, E., Regalado-Santiago, C., Millán-Aldaco, D., Blanco-Centurión, C., Drucker-Colín, R., 2011. Growth hormone improves hippocampal adult cell survival and counteracts the inhibitory effect of prolonged sleep deprivation on cell proliferation. Brain Res. Bull. 84, 252–257. 10.1016/j.brainresbull.2011.01.003

Gratwicke, J., Kahan, J., Zrinzo, L., Hariz, M., Limousin, P., Foltynie, T., Jahanshahi, M., 2013. The nucleus basalis of Meynert: A new target for deep brain stimulation in dementia? Neurosci. Biobehav. Rev. 37, 2676–2688. 10.1016/j.neubiorev.2013.09.003

Gratwicke, J., Zrinzo, L., Kahan, J., Peters, A., Beigi, M., Akram, H., Hyam, J., Oswal, A., Day, B., Mancini, L., Thornton, J., Yousry, T., Limousin, P., Hariz, M., Jahanshahi, M., Foltynie, T., 2018. Bilateral Deep Brain Stimulation of the Nucleus Basalis of Meynert for Parkinson Disease Dementia: A Randomized Clinical Trial. JAMA Neurol. 75, 169–178. 10.1001/jamaneurol.2017.3762

Gratwicke, J., Zrinzo, L., Kahan, J., Peters, A., Brechany, U., McNichol, A., Beigi, M., Akram, H., Hyam, J., Oswal, A., Day, B., Mancini, L., Thornton, J., Yousry, T., Crutch, S.J., Taylor, J.-P., McKeith, I., Rochester, L., Schott, J.M., Limousin, P., Burn, D., Rossor, M.N., Hariz, M., Jahanshahi, M., Foltynie, T., 2020. Bilateral nucleus basalis of Meynert deep brain stimulation for dementia with Lewy bodies: A randomised clinical trial. Brain Stimulat. 13, 1031–1039. 10.1016/j.brs.2020.04.010

Guzman-Marin, R., Bashir, T., Suntsova, N., Szymusiak, R., McGinty, D., 2007. Hippocampal neurogenesis is reduced by sleep fragmentation in the adult rat. Neuroscience 148, 325–333. 10.1016/j.neuroscience.2007.05.030

Hairston, I.S., Little, M.T.M., Scanlon, M.D., Barakat, M.T., Palmer, T.D., Sapolsky, R.M., Heller, H.C., 2005. Sleep restriction suppresses neurogenesis induced by hippocampus-dependent learning. J. Neurophysiol. 94, 4224–4233. 10.1152/jn.00218.2005

Han, Y., Shi, Y., Xi, W., Zhou, R., Tan, Z., Wang, H., Li, X., Chen, Z., Feng, G., Luo, M., Huang, Z., Duan, S., Yu, Y., 2014. Selective activation of cholinergic basal forebrain neurons induces immediate sleep-wake transitions. Curr. Biol. CB 24, 693–698. 10.1016/j.cub.2014.02.011

Hardenacke, K., Hashemiyoon, R., Visser-Vandewalle, V., Zapf, A., Freund, H.J., Sturm, V., Hellmich, M., Kuhn, J., 2016. Deep Brain Stimulation of the Nucleus Basalis of Meynert in Alzheimer’s Dementia: Potential Predictors of Cognitive Change and Results of a Long-Term Follow-Up in Eight Patients. Brain Stimulat. 9, 799–800. 10.1016/j.brs.2016.05.013

Headley, D.B., Paré, D., 2017. Common oscillatory mechanisms across multiple memory systems. Npj Sci. Learn. 2, 1. 10.1038/s41539-016-0001-2

Huang, C., Chu, H., Ma, Y., Zhou, Z., Dai, C., Huang, X., Fang, L., Ao, Q., Huang, D., 2019. The neuroprotective effect of deep brain stimulation at nucleus basalis of Meynert in transgenic mice with Alzheimer’s disease. Brain Stimulat. 12, 161–174. 10.1016/j.brs.2018.08.015

Irmak, S.O., de Lecea, L., 2014. Basal forebrain cholinergic modulation of sleep transitions. Sleep 37, 1941–1951. 10.5665/sleep.4246

Jack, C.R., Knopman, D.S., Jagust, W.J., Petersen, R.C., Weiner, M.W., Aisen, P.S., Shaw, L.M., Vemuri, P., Wiste, H.J., Weigand, S.D., Lesnick, T.G., Pankratz, V.S., Donohue, M.C., Trojanowski, J.Q., 2013. Tracking pathophysiological processes in Alzheimer’s disease: an updated hypothetical model of dynamic biomarkers. Lancet Neurol. 12, 207–216. 10.1016/S1474-4422(12)70291-0

Ju, Y.-E.S., McLeland, J.S., Toedebusch, C.D., Xiong, C., Fagan, A.M., Duntley, S.P., Morris, J.C., Holtzman, D.M., 2013. Sleep quality and preclinical Alzheimer disease. JAMA Neurol. 70, 587–593. 10.1001/jamaneurol.2013.2334

Junek, A., Rusak, B., Semba, K., 2010. Short-term sleep deprivation may alter the dynamics of hippocampal cell proliferation in adult rats. Neuroscience 170, 1140–1152. 10.1016/j.neuroscience.2010.08.018

Kalinchuk, A.V., McCarley, R.W., Stenberg, D., Porkka-Heiskanen, T., Basheer, R., 2008. The role of cholinergic basal forebrain neurons in adenosine-mediated homeostatic control of sleep: lessons from 192 IgG-saporin lesions. Neuroscience 157, 238–253. 10.1016/j.neuroscience.2008.08.040

Kang, J.-E., Lim, M.M., Bateman, R.J., Lee, J.J., Smyth, L.P., Cirrito, J.R., Fujiki, N., Nishino, S., Holtzman, D.M., 2009. Amyloid-beta dynamics are regulated by orexin and the sleep-wake cycle. Science 326, 1005–1007. 10.1126/science.1180962

Kuhn, J., Hardenacke, K., Lenartz, D., Gruendler, T., Ullsperger, M., Bartsch, C., Mai, J.K., Zilles, K., Bauer, A., Matusch, A., Schulz, R.-J., Noreik, M., Bührle, C.P., Maintz, D., Woopen, C., Häussermann, P., Hellmich, M., Klosterkötter, J., Wiltfang, J., Maarouf, M., Freund, H.-J., Sturm, V., 2015a. Deep brain stimulation of the nucleus basalis of Meynert in Alzheimer’s dementia. Mol. Psychiatry 20, 353–360. 10.1038/mp.2014.32

Kuhn, J., Hardenacke, K., Shubina, E., Lenartz, D., Visser-Vandewalle, V., Zilles, K., Sturm, V., Freund, H.J., 2015b. Deep Brain Stimulation of the Nucleus Basalis of Meynert in Early Stage of Alzheimer’s Dementia. Brain Stimulat. 8, 838–839. 10.1016/j.brs.2015.04.002

Kumbhare, D., Palys, V., Toms, J., Wickramasinghe, C.S., Amarasinghe, K., Manic, M., Hughes, E., Holloway, K.L., 2018. Nucleus Basalis of Meynert Stimulation for Dementia: Theoretical and Technical Considerations. Front. Neurosci. 12, 614. 10.3389/fnins.2018.00614

Kurosawa, M., Sato, A., Sato, Y., 1989. Stimulation of the nucleus basalis of Meynert increases acetylcholine release in the cerebral cortex in rats. Neurosci. Lett. 98, 45–50. 10.1016/0304-3940(89)90371-6

Lee, D.J., Milosevic, L., Gramer, R., Sasikumar, S., Al-Ozzi, T.M., Vloo, P.D., Dallapiazza, R.F., Elias, G.J.B., Cohn, M., Kalia, S.K., Hutchison, W.D., Fasano, A., Lozano, A.M., 2019. Nucleus basalis of Meynert neuronal activity in Parkinson’s disease. J. Neurosurg. 132, 574–582. 10.3171/2018.11.JNS182386

Linster, C., Garcia, P.A., Hasselmo, M.E., Baxter, M.G., 2001. Selective loss of cholinergic neurons projecting to the olfactory system increases perceptual generalization between similar, but not dissimilar, odorants. Behav. Neurosci. 115, 826–833. 10.1037//0735-7044.115.4.826

Liu, H., Wolters, A., Temel, Y., Alosaimi, F., Jahanshahi, A., Hescham, S., 2022. Deep brain stimulation of the nucleus basalis of Meynert in an experimental rat model of dementia: Stimulation parameters and mechanisms. Neurobiol. Dis. 171, 105797. 10.1016/j.nbd.2022.105797

Liu, R., Crawford, J., Callahan, P.M., Terry, A.V., Constantinidis, C., Blake, D.T., 2017. Intermittent Stimulation of the Nucleus Basalis of Meynert Improves Working Memory in Adult Monkeys. Curr. Biol. CB 27, 2640–2646.e4. 10.1016/j.cub.2017.07.021

Maltête, D., Wallon, D., Bourilhon, J., Lefaucheur, R., Danaila, T., Thobois, S., Defebvre, L., Dujardin, K., Houeto, J.-L., Godefroy, O., Krystkowiak, P., Martinaud, O., Gillibert, A., Chastan, M., Vera, P., Hannequin, D., Welter, M.-L., Derrey, S., 2020. Nucleus basalis of meynert stimulation for lewy body dementia: A phase I randomized clinical trial. Neurology 10.1212/WNL.0000000000011227. 10.1212/WNL.0000000000011227

Mann, A., Gondard, E., Tampellini, D., Milsted, J.A.T., Marillac, D., Hamani, C., Kalia, S.K., Lozano, A.M., 2018. Chronic deep brain stimulation in an Alzheimer’s disease mouse model enhances memory and reduces pathological hallmarks. Brain Stimulat. 11, 435–444. 10.1016/j.brs.2017.11.012

Mirescu, C., Peters, J.D., Noiman, L., Gould, E., 2006. Sleep deprivation inhibits adult neurogenesis in the hippocampus by elevating glucocorticoids. Proc. Natl. Acad. Sci. U. S. A. 103, 19170–19175. 10.1073/pnas.0608644103

Nombela, C., Lozano, A., Villanueva, C., Barcia, J.A., 2019. Simultaneous Stimulation of the Globus Pallidus Interna and the Nucleus Basalis of Meynert in the Parkinson-Dementia Syndrome. Dement. Geriatr. Cogn. Disord. 47, 19–28. 10.1159/000493094

Pagonabarraga, J., Kulisevsky, J., 2012. Cognitive impairment and dementia in Parkinson’s disease. Neurobiol. Dis. 46, 590–596. 10.1016/j.nbd.2012.03.029

Paxinos, G., Watson, C., n.d. The Rat Brain in Stereotaxic Coordinates - 6th Edition [WWW Document]. URL https://www.elsevier.com/books/the-rat-brain-in-stereotaxic-coordinates/paxinos/978-0-12-374121-9 (accessed 4.27.21).

Perry, E.K., Curtis, M., Dick, D.J., Candy, J.M., Atack, J.R., Bloxham, C.A., Blessed, G., Fairbairn, A., Tomlinson, B.E., Perry, R.H., 1985. Cholinergic correlates of cognitive impairment in Parkinson’s disease: comparisons with Alzheimer’s disease. J. Neurol. Neurosurg. Psychiatry 48, 413–421. 10.1136/jnnp.48.5.413

Roman, V., Van der Borght, K., Leemburg, S.A., Van der Zee, E.A., Meerlo, P., 2005. Sleep restriction by forced activity reduces hippocampal cell proliferation. Brain Res. 1065, 53–59. 10.1016/j.brainres.2005.10.020

Rothman, S.M., Herdener, N., Frankola, K.A., Mughal, M.R., Mattson, M.P., 2013. Chronic mild sleep restriction accentuates contextual memory impairments, and accumulations of cortical Aβ and pTau in a mouse model of Alzheimer’s Disease. Brain Res. 1529, 200–208. 10.1016/j.brainres.2013.07.010

Sarter, M., Parikh, V., Howe, W.M., 2009. Phasic acetylcholine release and the volume transmission hypothesis: time to move on. Nat. Rev. Neurosci. 10, 383–390. 10.1038/nm2635

Scarmeas, N., Brandt, J., Blacker, D., Albert, M., Hadjigeorgiou, G., Dubois, B., Devanand, D., Honig, L., Stern, Y., 2007. Disruptive behavior as a predictor in Alzheimer disease. Arch. Neurol. 64, 1755–1761. 10.1001/archneur.64.12.1755

Sims, J.R., Zimmer, J.A., Evans, C.D., Lu, M., Ardayfio, P., Sparks, J., Wessels, A.M., Shcherbinin, S., Wang, H., Monkul Nery, E.S., Collins, E.C., Solomon, P., Salloway, S., Apostolova, L.G., Hansson, O., Ritchie, C., Brooks, D.A., Mintun, M., Skovronsky, D.M., TRAILBLAZER-ALZ 2 Investigators, 2023. Donanemab in Early Symptomatic Alzheimer Disease: The TRAILBLAZER-ALZ 2 Randomized Clinical Trial. JAMA 330, 512–527. 10.1001/jama.2023.13239

Sportiche, N., Suntsova, N., Methippara, M., Bashir, T., Mitrani, B., Szymusiak, R., McGinty, D., 2010. Sustained sleep fragmentation results in delayed changes in hippocampal-dependent cognitive function associated with reduced dentate gyrus neurogenesis. Neuroscience 170, 247–258. 10.1016/j.neuroscience.2010.06.038

Stone, S.S.D., Teixeira, C.M., Devito, L.M., Zaslavsky, K., Josselyn, S.A., Lozano, A.M., Frankland, P.W., 2011. Stimulation of entorhinal cortex promotes adult neurogenesis and facilitates spatial memory. J. Neurosci. Off. J. Soc. Neurosci. 31, 13469–13484. 10.1523/JNEUROSCI.3100-11.2011

Suga, N., 2020. Plasticity of the adult auditory system based on corticocortical and corticofugal modulations. Neurosci. Biobehav. Rev. 113, 461–478. 10.1016/j.neubiorev.2020.03.021

Teipel, S.J., Temp, A.G.M., Lutz, M.W., 2024. Bayesian meta-analysis of phase 3 results of aducanumab, lecanemab, donanemab, and high-dose gantenerumab in prodromal and mild Alzheimer’s disease. Alzheimers Dement. Transl. Res. Clin. Interv. 10, e12454. 10.1002/trc2.12454

Unal, C., Golowasch, J., Zaborszky, L., 2012. Adult mouse basal forebrain harbors two distinct cholinergic populations defined by their electrophysiology. Front. Behav. Neurosci. 6.

Warm, J.S., Parasuraman, R., Matthews, G., 2008. Vigilance Requires Hard Mental Work and Is Stressful. Hum. Factors 50, 433–441. 10.1518/001872008X312152

Weinberger, N.M., Miasnikov, A.A., Bieszczad, K.M., Chen, J.C., 2013. Gamma band plasticity in sensory cortex is a signature of the strongest memory rather than memory of the training stimulus. Neurobiol. Learn. Mem. 104, 49–63. 10.1016/j.nlm.2013.05.001

Weinberger, N.M., Miasnikov, A.A., Chen, J.C., 2009. Sensory memory consolidation observed: increased specificity of detail over days. Neurobiol. Learn. Mem. 91, 273–286. 10.1016/j.nlm.2008.10.012

Weinberger, N.M., Miasnikov, A.A., Chen, J.C., 2006. The level of cholinergic nucleus basalis activation controls the specificity of auditory associative memory. Neurobiol. Learn. Mem. 86, 270–285. 10.1016/j.nlm.2006.04.004

Yi, F., Catudio-Garrett, E., Gábriel, R., Wilhelm, M., Erdelyi, F., Szabo, G., Deisseroth, K., Lawrence, J., 2015. Hippocampal “cholinergic interneurons” visualized with the choline acetyltransferase promoter: anatomical distribution, intrinsic membrane properties, neurochemical characteristics, and capacity for cholinergic modulation. Front. Synaptic Neurosci. 7. 10.3389/fnsyn.2015.00004

Zant, J.C., Kim, T., Prokai, L., Szarka, S., McNally, J., McKenna, J.T., Shukla, C., Yang, C., Kalinchuk, A.V., McCarley, R.W., Brown, R.E., Basheer, R., 2016. Cholinergic Neurons in the Basal Forebrain Promote Wakefulness by Actions on Neighboring Non-Cholinergic Neurons: An Opto-Dialysis Study. J. Neurosci. Off. J. Soc. Neurosci. 36, 2057–2067. 10.1523/JNEUROSCI.3318-15.2016

